# RAB7 deficiency impairs pulmonary artery endothelial function and promotes pulmonary hypertension

**DOI:** 10.1101/2023.02.03.526842

**Authors:** Bryce Piper, Srimathi Bogamuwa, Tanvir Hossain, Daniela Farkas, Lorena Rosas, Adam Green, Geoffrey Newcomb, Nuo Sun, Jeffrey C. Horowitz, Aneel R Bhagwani, Hu Yang, Tatiana V. Kudryashova, Mauricio Rojas, Ana L. Mora, Pearlly Yan, Rama K. Mallampalli, Elena A. Goncharova, David M. Eckmann, Laszlo Farkas

## Abstract

Pulmonary arterial hypertension (PAH) is a devastating and progressive disease with limited treatment options. Endothelial dysfunction plays a central role in development and progression of PAH, yet the underlying mechanisms are incompletely understood. The endosome-lysosome system is important to maintain cellular health and the small GTPase RAB7 regulates many functions of this system. Here, we explored the role of RAB7 in endothelial cell (EC) function and lung vascular homeostasis. We found reduced expression of RAB7 in ECs from PAH patients. Endothelial haploinsufficiency of RAB7 caused spontaneous PH in mice. Silencing of RAB7 in ECs induced broad changes in gene expression revealed via RNA sequencing and RAB7 silenced ECs showed impaired angiogenesis, expansion of a senescent cell fraction, combined with impaired endolysosomal trafficking and degradation, which suggests inhibition of autophagy at the pre-degradation level. Further, mitochondrial membrane potential and oxidative phosphorylation were decreased, and glycolysis was enhanced. Treatment with the RAB7 activator ML-098 reduced established PH in chronic hypoxia/SU5416 rats. In conclusion, we demonstrate here for the first time the fundamental impairment of EC function by loss of RAB7 that leads to PH and show RAB7 activation as a potential therapeutic strategy in a preclinical model of PH.

## INTRODUCTION

Pulmonary arterial hypertension (PAH) is a severe and fatal condition characterized by increased pulmonary artery pressure and extensive remodeling of all layers of the pulmonary artery wall (1, 2). The molecular pathogenesis of PAH remains incompletely understood at this time (1). One important and defining feature of pulmonary arterial dysfunction and remodeling in PAH is lung endothelial cell (EC) dysfunction. Following endothelial injury and apoptosis, endothelial dysfunction presents as unchecked proliferation likely following endothelial injury and apoptosis, impaired angiogenesis, enhanced release of pro-inflammatory cytokines and upregulation of adhesion molecules (1). Similar to cancer, the hyperproliferative ECs in PAH may derive from clonal expansion of primitive stem-cell like ECs (3, 4). Our previous work indicates also that while these primitive, hyperproliferative ECs can promote lung vascular remodeling, this remodeling is reversible after removal of additional triggers, such as hypoxia (4). This result indicates that proliferation alone may not be sufficient to develop progressive pulmonary vascular remodeling and PAH. Recently, a subpopulation of senescent ECs has been identified in PAECs from patients with several forms of pulmonary hypertension (PH), including PAH, and these senescent ECs appear to promote the irreversibility of PA remodeling and PH (5, 6). We will use the term “PH” when describing animal models throughout the manuscript. The dysfunctional ECs from PAH patients further exhibit dysfunction of mitochondrial function, including dysregulation of mitochondrial membrane potential ΔΨ_m_, reduced mitochondrial mass, impaired oxidative phosphorylation and glycolytic shift akin to cancer cells (7–10). PAH ECs show reduced oxidative phosphorylation, but the impaired regulation of mitochondrial health in PAH ECs is not sufficiently explained (8). Key cellular functions are regulated by autophagy, or the endolysosomal degradation of macromolecules and cell organelles, including mitochondria (11). In animal models of PH, conflicting results either attribute a protective function to physiologic autophagy or indicate that inhibition of aberrant autophagy is a treatment for PH (12, 13). Endosomal sorting, trafficking and fusion of endosomes and lysosomes to endolysosomes are critical steps in physiologic autophagy (14). The small GTPase RAB7A has a key function in all these steps (15, 16). RAB7A is commonly referred to as RAB7, which we will use throughout the manuscript. Consequently, loss of RAB7 disrupts cellular function and impairs mitochondrial health, amplifying tissue injury (17–20).

We hypothesized that endothelial RAB7 deficiency is a cause of endothelial dysfunction in PAH through inhibition of autophagy on the pre-degradation level and promotes lung vascular remodeling and PH. We show here for the first time reduced RAB7 expression in pulmonary artery ECs (PAECs) from PAH patients and identified that endothelial-specific reduction of RAB7 expression caused spontaneous PH in mice. *In vitro*, RAB7 knockdown impaired endolysosomal trafficking and degradation and fundamentally altered gene expression in PAECs with functional evidence of impaired angiogenesis and expansion of a senescent EC subpopulation. In RAB7 deficient ECs, we detected the loss of ΔΨ_m_ with a shift towards dysfunctional mitochondria and evidence of reduced mitochondrial respiration with enhanced glycolysis. The RAB7 GTPase activator ML-098 reduced severe PH in Hx/Su rats, hence supporting RAB7 as a potential therapeutic target in PAH.

## RESULTS

### Endosomal GTPase RAB7 is reduced in PAECs from patients with PAH

We first identified the expression of RAB7 in lung tissues, PAECs and PASMCs from patients with PAH. The expression of RAB7 was reduced in vWF^+^ PAECs in concentric and plexiform lesions from PAH patients compared to control lungs (**Fig. 1A**). PAH PAECs had reduced expression of RAB7, whereas PAH PASMCs had similar RAB7 protein expression levels as control PAEC (**Fig. 1B, C**).

**Figure 1.**
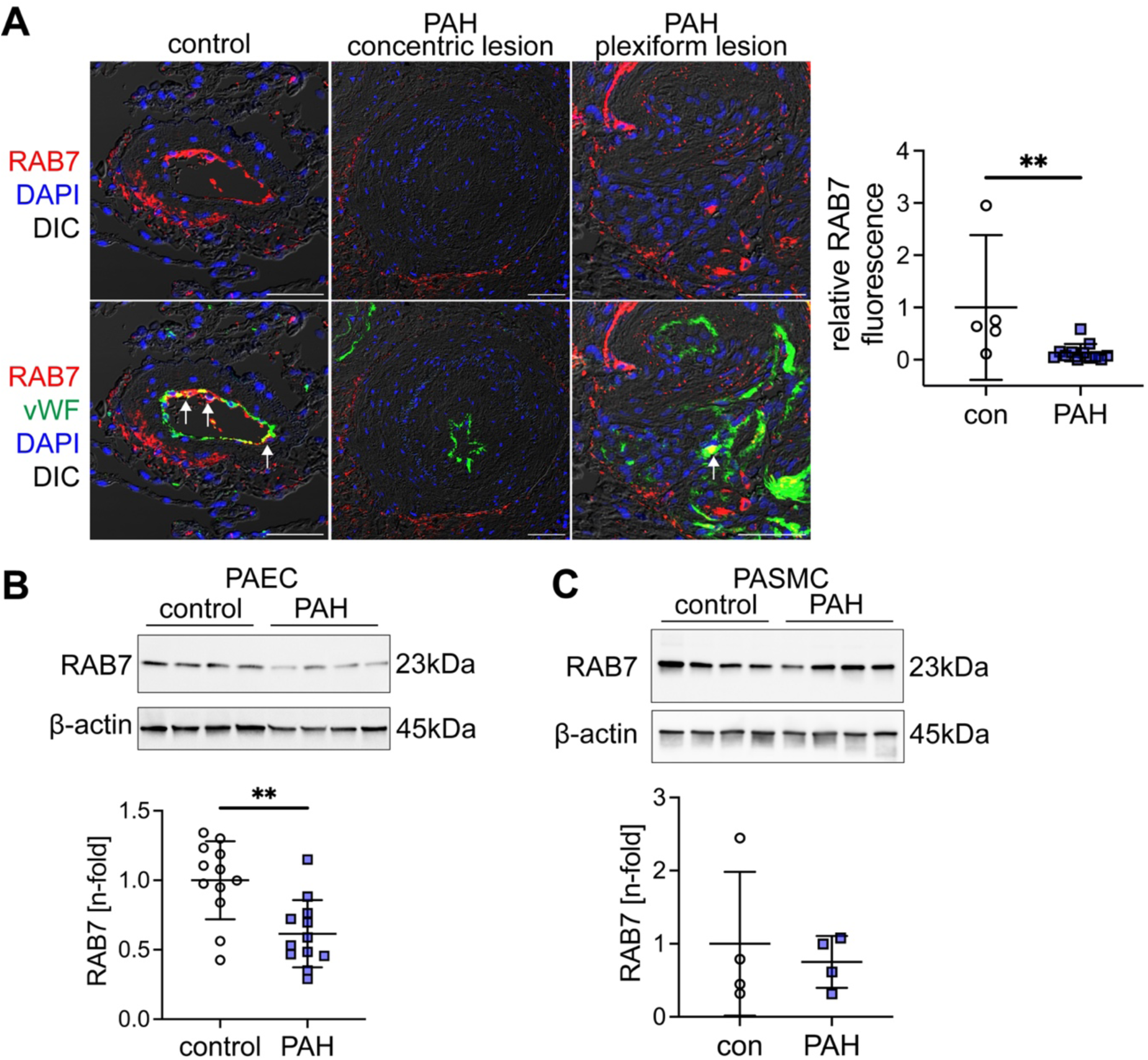
Reduced RAB7 expression in endothelial cells from PAH patients. (**A**) Representative optical sections (confocal microscopy, representative of n=3 patients per group) show reduced RAB7 expression in ECs in concentric and plexiform lesions from PAH patients compared to controls. Arrows indicate vWF^+^ ECs with strong RAB7 expression. Scale bar: 50μm. Nuclear staining with DAPI. Graph shows quantification of relative RAB7 immunofluorescence in ECs from control and PAH pulmonary arteries. (**B-C**) Representative Immunoblot and quantification of RAB7 in PAECs (B) from n=12 control (failed donor) and n=12 PAH patients. (C) shows representative immunoblot and quantification of RAB7 in PASMCs from n=4 control (failed donor) and n=4 PAH patients. β-actin is shown as loading control. All graphs show single values and mean±SD. **P<0.01.

### Endothelial RAB7 haploinsufficiency causes spontaneous PH

We then studied RAB7 expression in the chronic hypoxia/SU5416 (Hx/Su) rat model of PH, because this model reproduces many features of PAH, including the occlusive pulmonary arteriopathy that is characteristic of PAH (21, 22). We found using lung tissue sections and protein lysate an overall reduction of RAB7 expression in the lung tissue of Hx/Su rats at day 21 and 42 and localized the reduction to the ECs in the remodeled pulmonary arteries of Hx/Su rats (**Fig. 2A-B**). Then, we generated heterozygous EC-specific RAB7 knockout (KO, endothelial RAB7 haploinsufficient) mice by crossbreeding RAB7^fl/fl^ mice with Cdh5-Cre mice. We found increased media wall thickness (MWT) and right ventricular systolic pressure (RVSP) and a decreased ratio of pulmonary artery acceleration time /pulmonary ejection time (PAT/PET) in RAB7^fl/wt^ Cdh5-Cre^+^ mice after exposure to Hx/Su, indicating exaggerated pulmonary artery remodeling and PH with endothelial RAB7 haploinsufficiency (**Fig. 2C-G**). Echocardiographically estimated right ventricular (RV) cardiac output was reduced in Hx/Su-exposed RAB7^fl/wt^ Cdh5-Cre^-^ mice without a significant difference between Cre^-^ and Cre^+^ mice (**Fig. 2F**).

**Figure 2.**
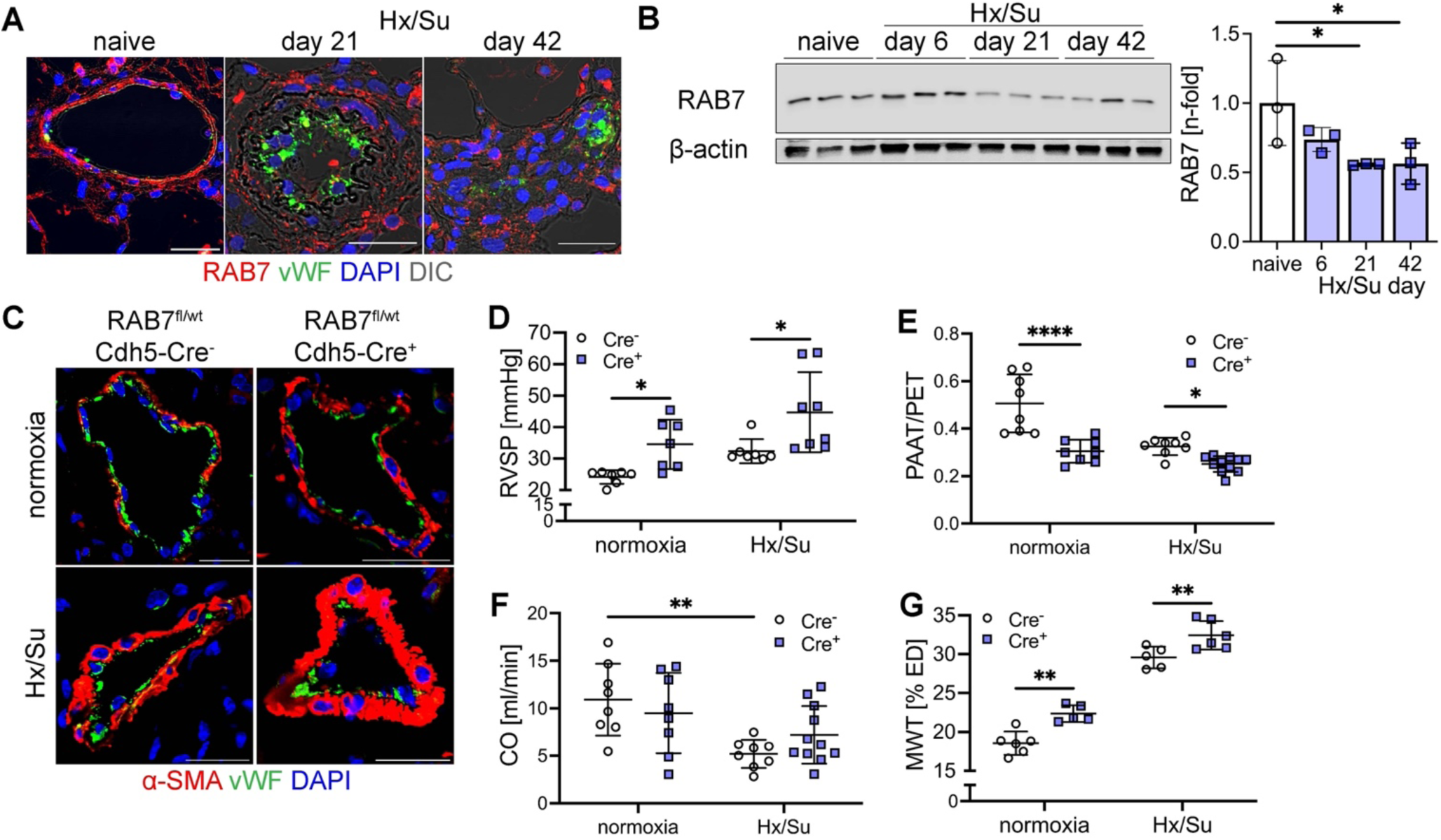
Loss of RAB7 expression causes PH in vivo. (**A**) Representative immunofluorescence images (of n=3) (confocal microscopy) show strong RAB7 staining in pulmonary arteries (PAs) from naïve rats, including in PAECs. In chronic hypoxia + SU5416 (Hx/Su) treated rats, RAB7 expression was particularly decreased in ECs in the remodeled PAs at day 21 and 42. Scale bars: 25 μm. (**B**) Representative immunoblot and densitometry of RAB7 expression in naïve and Hx/Su rats. (**C**) Representative double immunofluorescence for von Willebrand Factor (vWF) and α-smooth muscle actin (α-SMA) (optical section, confocal microscopy). Scale bars: 25 μm. (**D-G**) Right ventricular systolic pressure (RVSP, D), pulmonary artery acceleration time vs. pulmonary ejection time ratio (PAT/PET, E), echocardiographically estimated cardiac output (CO, F) and media wall thickness (MWT, G) in pulmonary arteries of RAB7^fl/wt^ Cdh5-Cre^-^ and RAB7^fl/wt^ Cdh5-Cre^+^ mice exposed to normoxia and Hx/Su. All graphs show single values and mean±SD. *P<0.05, **P<0.01, ****P<0.0001.

### Gene silencing of RAB7 induces endothelial dysfunction in PAECs

We then tested if RAB7 expression contributes to physiological EC function. Using siRNA targeting of RAB7 and bulk RNA sequencing (RNAseq) we found that RAB7 knockdown resulted in differential expression of 4842 genes using analysis parameters of fold change >1.25 and adjusted P value <0.05. We detected a significant amount of differentially expressed genes (DEGs) with function in cell cycle and DNA repair, cellular movement and trafficking, immune function and inflammation, and development (**Fig. 3**). Genes affecting angiogenesis and EC barrier function were among the DEGs with the highest n-fold changes in RAB7 siRNA treated PAECs. We found upregulation of anti-angiogenic genes *PCDH17, IGFBP6* and *CH25H*, and downregulation of angiogenic and EC function genes *DLL4, UNC5B, CLDN5, RASIP1, PGF, GJA5, EPN3, CCM2L and TSPAN18* (**Fig. 4A**). We then tested EC function *in vitro* and found that knockdown of RAB7 inhibited angiogenic network formation and gap closure, indicating impaired migration (**Fig. 4B-C**). Consistent with these functional changes our RNAseq data revealed a pro-senescent transcriptomic signature in RAB7 siRNA treated PAECs (**Fig. 4D**). These findings were mirrored by elevated p16 protein level (**Fig. 4E**) and increased fraction of senescence-associated β-galactosidase (SA-β-gal)^+^ PAECs (**Fig. 4F**).

**Figure 3.**
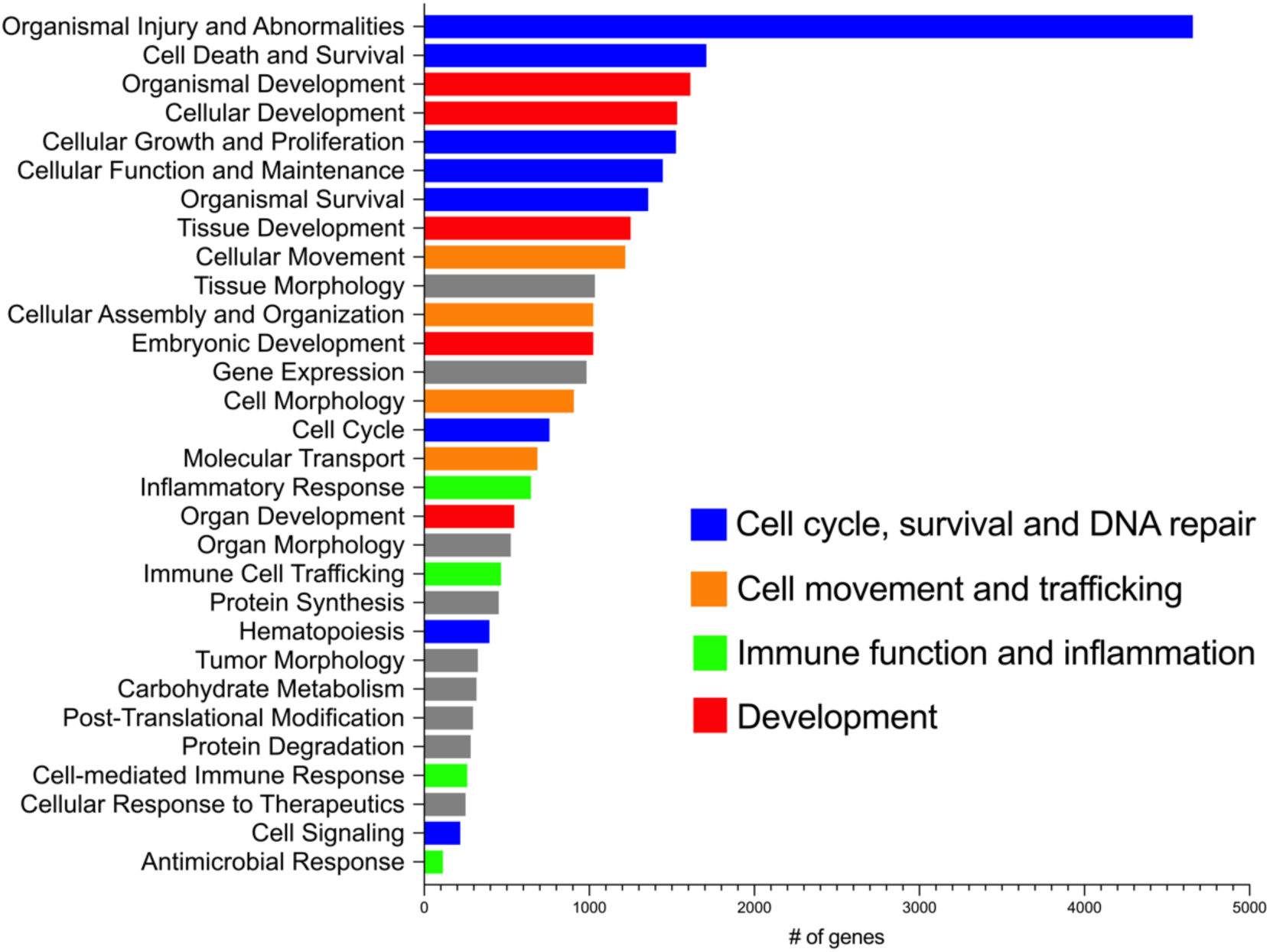
RNA sequencing of RAB7 silenced pulmonary artery ECs. Ingenuity pathway analysis of RNA sequencing data in PAECs transfected with RAB7 siRNA vs. control siRNA (fold change >1.25, adjusted P value <0.05). Diagram shows the 30 most regulated function terms showing the number of differentially expressed genes (DEG) in the RNAseq dataset within each category. DEG terms were labeled according to their relevance for cell cycle and DNA repair (blue), cellular movement and trafficking (orange), immune function and inflammation (green) and development (red).

**Figure 4.**
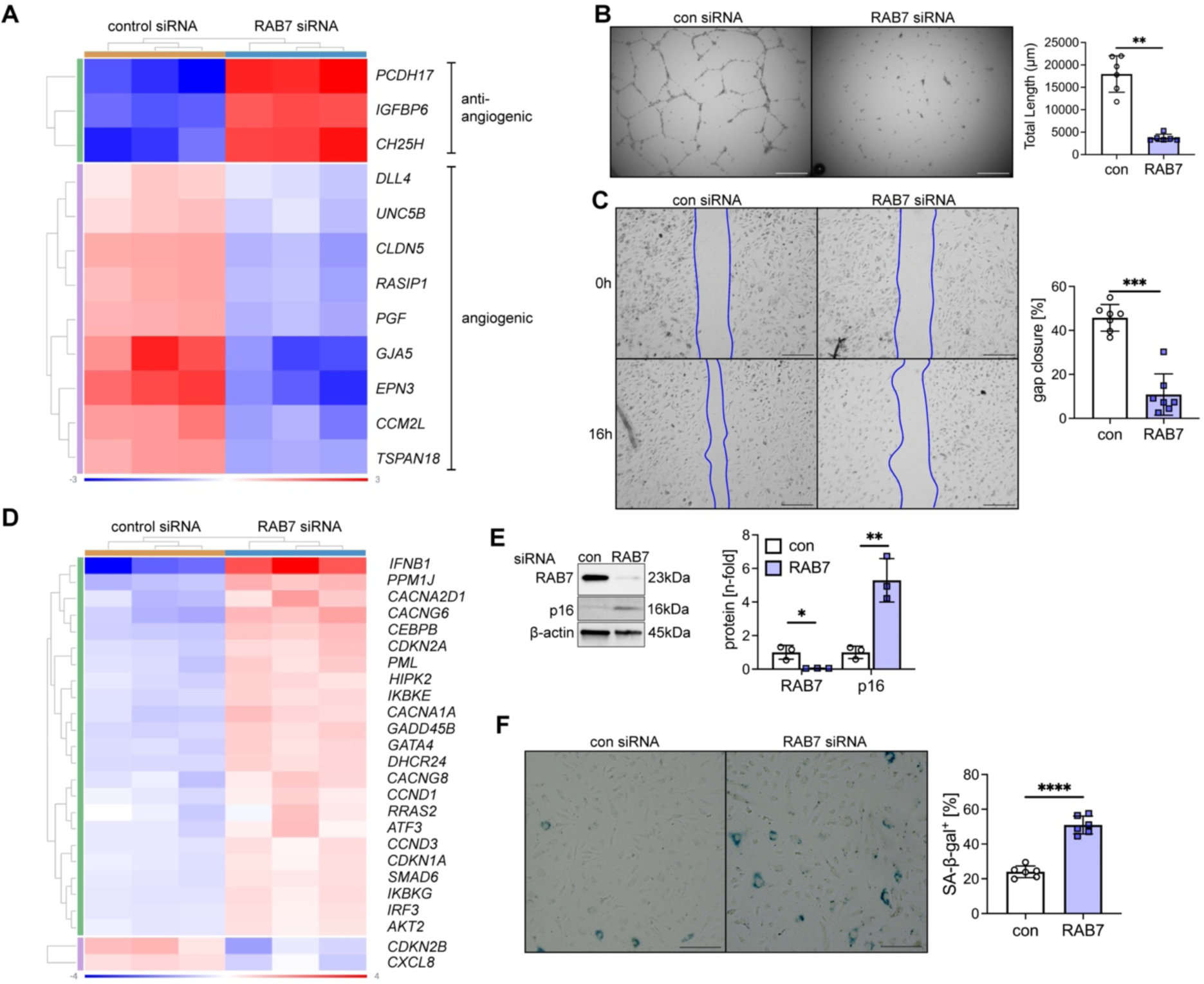
Loss of RAB7 induces endothelial dysfunction and senescence in pulmonary artery ECs. (**A**) Clustered heatmap demonstrates increased expression of anti-angiogenic genes, and reduced expression of angiogenic and EC barrier function genes in RNAseq of PAECs after RAB7 siRNA treatment vs. control siRNA. (**B**) Representative phase contrast images after 24h and quantification of total network length in RAB7 siRNA treated PAECs. n=6 per group. (**C**) Representative phase contrast after 16h of gap closure assay and quantification of percent gap closure in RAB7 siRNA transfected PAECs vs. control siRNA. n=7 per group. (**D**) Clustered heatmap of DEGs of senescence-associated gene expression pattern (Ingenuity Pathway Analysis) in RNAseq of PAECs treated with RAB7 siRNA. (**E**) Representative immunoblot and densitometric quantification for RAB7 and p16 in PAECs treated with RAB7 siRNA. n=3 per group. (**F**) Representative images and quantification of the fraction of SA-β-gal^+^ PAECs after RAB7 siRNA treatment vs. control siRNA. n=6 per group. Data are shown as single values and mean±SD. Heatmap data is normalized log_2_ fold expression. Scale bars: 200 μm (F), 250 μm (B, C). *P<0.05, **P<0.01, ***P<0.001, ****P<0.0001.

### RAB7 is required for endolysosome function in PAECs

Endolysosome function requires RAB7 (15, 23–25). To identify the extent to which RAB7 is required for endolysosome function in PAECs, we studied intracellular endolysosome trafficking in PAECs from PAH patients and in control PAECs following RAB7 knockdown. Using pHrodo dextran, which emits fluorescence after acidification in endosome and lysosome, we detected that dextran accumulates in PAH PAECs in enlarged vesicles (**Fig. 5A**). Using baculovirus-mediated expression of the GFP-labeled RAB5A, a marker of the early endosome, we found that pHrodo dextran accumulates in enlarged RAB5A^+^ early endosomes following endosomal uptake in RAB7-silenced PAECs (**Fig. 5B**). RAB7 siRNA also caused the accumulation of pHrodo dextran in enlarged lysosomes which we detected by staining with lysotracker (**Fig. 5C**). Consistently, our RNAseq data revealed the downregulation of multiple autophagy-related genes with RAB7 siRNA (**Fig. 5D**). Reduced cathepsin B activity further supported reduced lysosomal autophagy in RAB7 silenced PAECs (**Fig. 5E**).

**Fig. 5.**
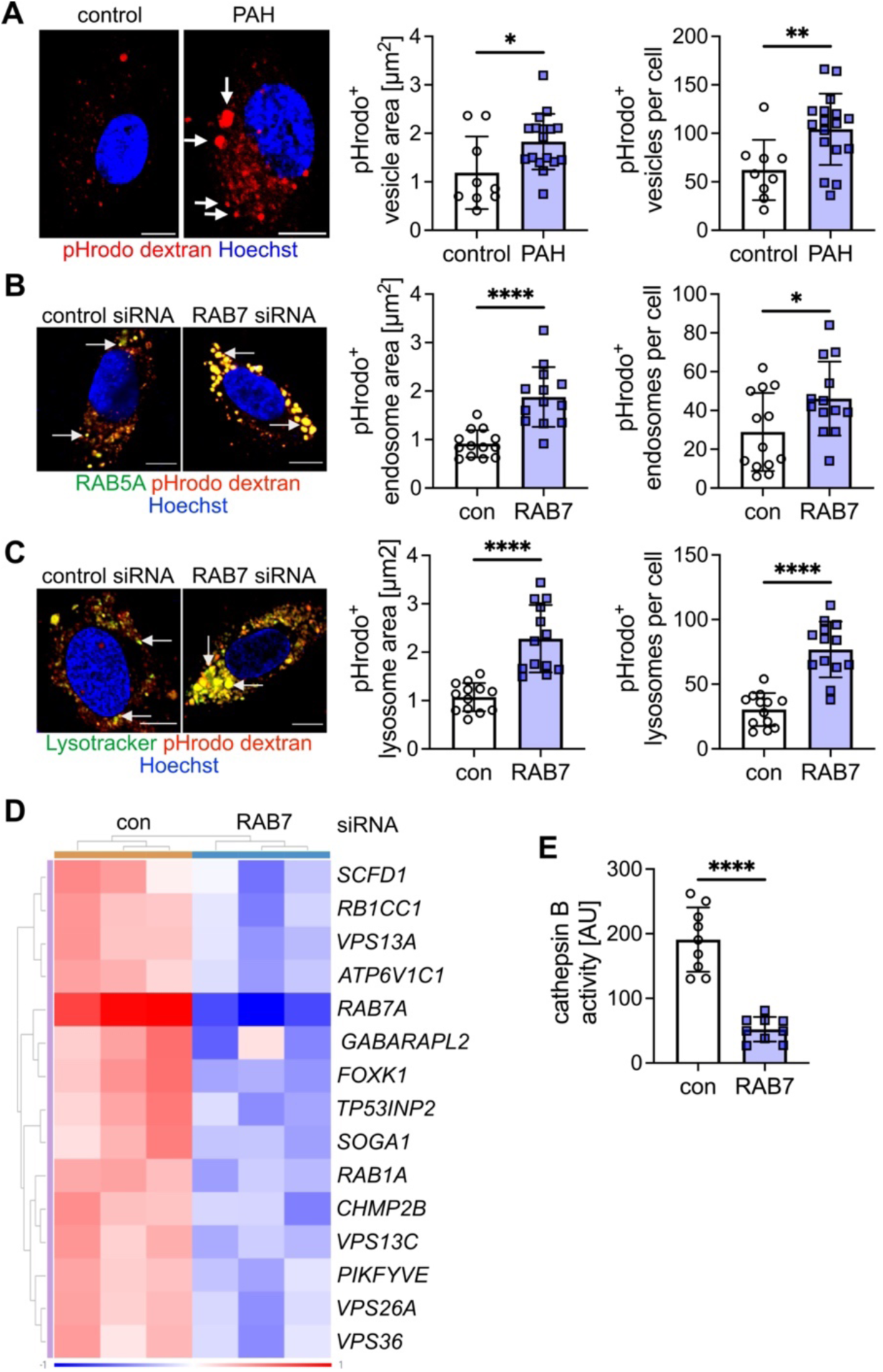
Impaired endosome-lysosome function in PAH pulmonary artery ECs and RAB7 deficient pulmonary artery ECs. (**A**) Representative optical sections (confocal microscopy) and quantification of pHrodo dextran^+^ vesicle area and number of vesicles per cell indicate accumulation of pHrodo dextran after 20 min in enlarged vesicles in PAH PAECs (arrows), but not in control PAECs. pHrodo dextran is taken up by endosomes and emits red fluorescence signal when pH drops during endosomal acidification. (**B**) Representative optical sections (confocal microscopy) show accumulation of pHrodo dextran after 20 min. in enlarged early endosomes. Early endosomes were identified by transfection of PAECs with baculovirus expressing GFP-labeled RAB5. Quantification of RAB5^+^ pHrodo dextran^+^ endosome area and number per cell confirms that dextran accumulates in enlarged early endosomes following RAB7 silencing. (**C**) Representative optical sections (confocal microscopy) show accumulation of pHrodo dextran after 20 min. in enlarged lysosomes. Lysosomes were labeled with lysotracker. Quantification of lysotracker^+^ pHrodo dextran^+^ lysosome area and number per cell confirms that dextran accumulates in enlarged lysosomes following RAB7 silencing. (**D**) Clustered heatmap shows autophagy-related DEG that were found to be downregulated in RNAseq from PAECs treated with RAB7 siRNA. Expression as normalized log_2_ fold. (**E**) Reduced cathepsin B activity also indicates impaired lysosomal autophagy. Scale bars: 10 μm. All graphs show single values and mean±SD. *P<0.05, **P<0.01, ****P<0.0001.

### RAB7 knockdown impairs mitochondrial membrane potential and oxidative phosphorylation

Because autophagy is important to maintain physiologic mitochondrial function (26), we tested the role of RAB7 in maintenance of the mitochondrial membrane potential ΔΨ_m_ and mitochondrial respiration. Using a flow cytometry assays for TMRE/mitotracker green, we found that RAB7 knockdown reduced the fraction of ECs with functional mitochondria (polarized, while increasing the fraction of ECs with dysfunctional (depolarized) mitochondria, indicating an overall reduction in ΔΨ_m_ (**Fig. 6A**). Using fluorescence microscopy of TMRM staining, RAB7 silencing caused a net reduction in ΔΨ_m_ with accumulation of the remaining functional mitochondria in the perinuclear region (**Fig. 6B**). We further detected increased mitochondrial net motility in perinuclear and peripheral regions in RAB7 siRNA PAECs (**Fig. 6B**). RAB7 silencing also increased mitochondrial production of reactive oxygen species (ROS) as measured by mitosox flow cytometry assay (**Fig. 6C**). RNAseq analysis of PAECs treated with RAB7 siRNA revealed upregulation of mRNA for multiple genes related to glycolysis, whereas the mRNA of various genes related to oxidative phosphorylation was reduced (**Fig. 6D**). RAB7 knockdown also impaired oxidative phosphorylation, because our data show reduced oxygen consumption rate (OCR) at baseline, maximal respiration, and spare respiratory capacity (**Fig. 6E**). There was no difference in proton leak. In addition, RAB7 siRNA treated PAECs had higher lactate production than control siRNA treated PAECs, indicating a shift towards glycolysis (**Fig. 6F**). Lastly, the extracellular acidification rate (ECAR) measurements revealed increased glycolysis, glycolytic capacity, and glycolytic reserve (**Fig. 6G**).

**Fig. 6.**
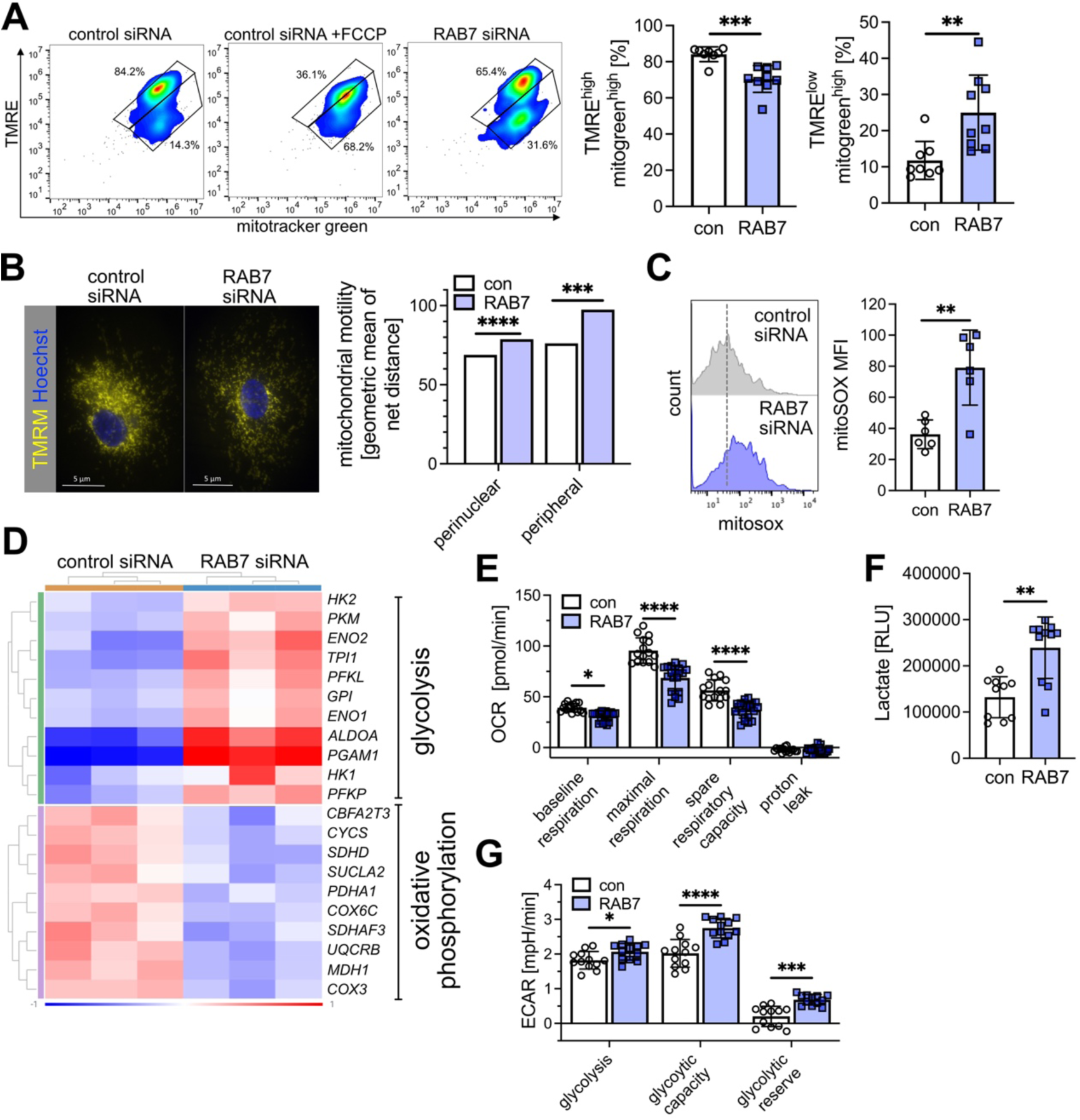
RAB7 silencing impairs mitochondrial membrane potential and mitochondrial function. (**A**) RAB7 siRNA impairs mitophagy as assessed by TMRE and mitotracker green (mitogreen) flow cytometry. FCCP is positive control (depolarizes mitochondrial membrane). TMRE^high^ mitogreen^high^ cells indicate cells with polarized mitochondrial membrane (functional mitochondria), whereas TMRE^low^ mitogreen^high^ cells are cells with depolarized mitochondrial membrane (dysfunctional mitochondria). N=8-9 per group. (**B**) Representative TMRM staining images show overall reduction and perinuclear accumulation of functional mitochondria in RAB7 siRNA versus control siRNA. Images are representative of n=3 experiments. Scale bar: 5 μm. RAB7 knockdown increases mitochondrial motility both in the peripheral and the perinuclear region. (**C**) RAB7 siRNA treatment promotes mitochondrial ROS production as indicated by flow cytometry for mitosox. n=6 per group. Figures shows representative histogram plots and quantification of mean fluorescence intensity (MFI). (**D**) Clustered heatmap shows glycolysis-related DEGs that were found to be upregulated and oxidative phosphorylation-related DEGs that were downregulated in RNAseq from PAECs treated with RAB7 siRNA. Expression as normalized log_2_ fold. (**E**) Seahorse high resolution respirometrics show reduced oxygen consumption rate (OCR) in RAB7 siRNA PAECs at basal respiration, maximal respiration and spare respiratory capacity. No difference was seen for proton leak. n=15 (con) and n=21 (RAB7). (**F**) Luminescence assay for lactate shows increased lactate production in RAB7 siRNA vs. control siRNA PAECs. n=10 (control) and n=11 (RAB7). (**G**) Extracellular acidification rate (ECAR) data show more rapid acidification (i.e., greater reliance on glycolysis) in the RAB7 siRNA PAECs as shown for glycolysis, basal and maximum glycolytic rate. n=12 per group. All graphs show single values and mean±SD (except F, as bar graphs in F indicate geometric mean of a log-normal distribution). *P<0.05, **P<0.01, ***P<0.001 and ****P<0.0001.

### RAB7 activator ML-098 reduces experimental PH in rats

To aid translation of our findings to a potential treatment option, we tested if the RAB7 GTPase activator ML-098 reduces experimental PH induced by Hx/Su in rats. First, we performed a preventive dose-response experiment to find the lowest efficacious dose. We found overall that 1.0 and 10 mg/kg doses reduced pulmonary artery occlusion, MWT and RVSP most effectively (**Fig. 7A-F**). Based on these results, we opted for 1.0 mg/kg in the interventional treatment approach. In this approach, ML-098 treatment reduced RVSP, Fulton index, MWT, pulmonary artery occlusion and mural cell proliferation while increasing PAT/PET and tricuspid annular plane systolic excursion (TAPSE) (**Fig. 7G-O**). We further found a trend towards elevated RV cardiac output with therapeutic ML-098 treatment (**Fig. 7P**). Hence, the RAB7 GTPase activator ML-098 prevents and reduces occlusive pulmonary artery occlusion, PH and RV dysfunction induced by Hx/Su.

**Figure 7.**
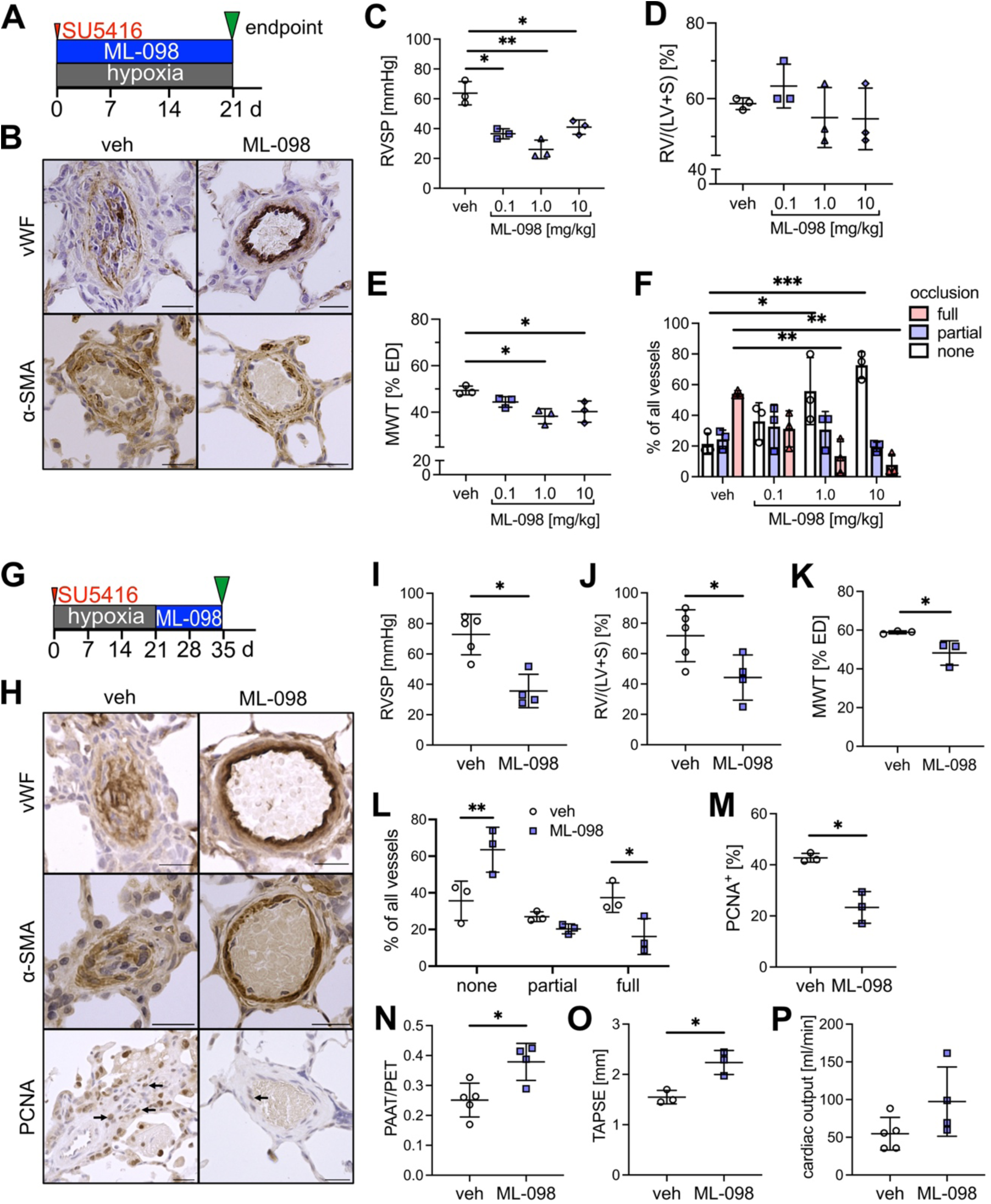
RAB7 activator ML-098 reduces PH in rats exposed to chronic hypoxia + SU5416. (**A**) Preventive treatment diagram. (**B**) Representative vWF and α-SMA immunohistochemistry images. (**C**) RVSP, (**D**) Fulton index, (**E**) pulmonary artery MWT and (**F**) occlusion of small pulmonary arteries. (**G**) interventional treatment diagram. (**H**) Representative vWF, α-SMA and PCNA immunohistochemistry. (**I**) RVSP, (**J**) Fulton index, (**K**) MWT of small pulmonary arteries, (**L**) occlusion of small pulmonary arteries, (**M**) percentage of PCNA^+^ pulmonary artery mural cells, (**N**) ratio of pulmonary artery acceleration time (PAAT) vs. pulmonary ejection time (PET), (**O**) tricuspid annular plain systolic excursion (TAPSE) and (**P**) echocardiographic estimation of cardiac output. Scale bars: 25 μm. *P<0.05, **P<0.01, ***P<0.001. n=3 (C-F, K, L, M, O), n=4-5 (I, J, N, P). All graphs show single values and mean±SD.

## DISCUSSION

PAH remains a deadly disease and current vasodilator therapies are not sufficient to cure the disease by a lacking effect on the progressive occlusive pulmonary arteriopathy that is a hallmark of PAH (1, 2). One potential underlying cause is that these treatments mainly improve the vasotonus function of ECs, but not the overall EC dysfunction (1, 2). As coordinated autophagy and mitochondrial function have been shown to be important drivers of physiologic EC function (7–11), we sought to find a unifying pathogenetic process that explains these findings. The endosome-lysosome system holds a key role in the trafficking, recycling, and degradation of macromolecules and cell organelles, such as mitochondria. The GTPase RAB7 is crucial for the regulation of endosome-lysosome function, autophagy, and cell function. Hence, we hypothesized that endothelial RAB7 deficiency is a cause of endothelial dysfunction, lung vascular remodeling and PAH.

The main findings of our study are that 1) PAECs from PAH patients have reduced RAB7 expression and impaired endosome-lysosome function; 2) endothelial haploinsufficiency of RAB7 induces spontaneous PH; 3) Silencing of RAB7 in human PAECs impairs endosome/lysosome function, autophagy and angiogenesis, and promotes cellular senescence; 4) RAB7 insufficiency reduced mitochondrial membrane potential ΔΨ_m_ and oxidative phosphorylation while promoting mitochondrial ROS production and glycolysis; 5) The RAB7 GTPase activator ML-098 reduced severe PH and right heart dysfunction induced by cHx/Su.

Whereas all layers of the pulmonary artery wall and all mural cell types are affected, EC dysfunction has emerged as one of the key drivers of pulmonary artery remodeling in PAH (27). ECs not only provide an important barrier, but in the pulmonary circulation also affect smooth muscle cell function by regulation of e.g., vasotonus or smooth muscle cell growth through release of mediators (27). In addition, altered EC function may contribute to lung vascular remodeling by microvascular dropout following EC apoptosis and to the development of complex plexiform lesions through aberrant cell growth (4, 28–30). Yet current therapeutic avenues are not successful in restoring physiologic EC function and reversing pulmonary artery remodeling.

Endosomes develop as plasma membrane invaginations during endocytosis and are a shuttle for macromolecules from the outside of the cell into the cell (31). In addition, internalization of receptors and other macromolecules from the cell membrane occurs also through the endosome system. After initial formation, endosomes are designated “early endosomes” and carry specific surface molecules, including early endosome antigen 1 (EEA1) and members of the RAB family of small GTPases, such as RAB5 (31, 32). During sorting, endosomal content including macromolecules and cell organelles is either being recycled to the cell surface or designated for degradation via autophagy. Upon maturation to multivesicular “late endosomes”, these fuse with lysosomes to endolysosomes as sites of autophagy (31). RAB7 is a GTPase with central function in endosomal sorting, trafficking and fusion with lysosomes (31). Not surprisingly, loss of RAB7 impairs autophagy and particularly autophagy of mitochondria, or mitophagy, and hence cell function (15, 18–20, 33–37). Our results show that PAECs, but not PASMCs, from patients with PAH have reduced expression of RAB7 and impaired endosomal trafficking and/or degradation of dextran. Consequently, endothelial RAB7 haploinsufficient mice developed spontaneous PH under normoxic conditions and more severe PH following exposure to Hx/Su, suggesting a fundamental protective role for physiologic levels of RAB7 in PAECs. Our results further show that RAB7 silencing in PAECs impairs endosomal and lysosomal trafficking and/or degradation, indicating that RAB7 is essential for autophagy. While knockout of the autophagy component Microtubule-associated proteins 1A/1B light chain 3B (LC3B) promoted PH in a mouse model and supports our findings, some studies have hinted at a potential role for excessive autophagy in EC dysfunction and PH progression (12, 13, 38, 39). A more detailed analysis of endosome/lysosome function and degradation processes in ECs and other vascular cell types will likely be required to close the gap towards a definitive answer.

Our focus here was on elucidating the functional role of RAB7 in PAECs and evidence is accumulating that mitochondrial dysfunction has a central role in the regulation of EC dysfunction and pulmonary artery remodeling (7, 9, 10, 40). RAB7 is a known modulator of mitochondrial health through mitochondrial fission and mitophagy, and we found that knockdown of RAB7 reduces mitochondrial membrane potential ΔΨ_m_ and impairs oxidative phosphorylation. These findings are consistent with enhanced glycolysis and lactate production, as well as impaired clearance of mitochondrial ROS. Impaired mitochondrial respiration and a shift towards glycolysis is a known feature of PAECs from PAH patients and are consistent with a dysregulation of EC function akin to a “Warburg effect” (7, 9, 10). Increased mitochondrial ROS has been previously shown in bone morphogenic protein receptor 2 (BMPR2) knockout ECs and reduced ΔΨ_m_ was caused in PAH PAECs by hypoxia-reoxygenation, but by normoxia exposure (9). Hence, our findings corroborate that RAB7 deficiency contributes to impaired mitochondrial health and mitochondria function in PAH PAECs.

Our results further show that silencing of RAB7 impairs angiogenesis and migration in PAECs, and promotes senescence in PAECs. Impaired angiogenesis has been previously shown in PAECs and late outgrowth endothelial progenitor cells from PAH patients (41, 42). While these studies focused on the proliferative phenotype of PAH PAECs, more recent studies have demonstrated that endothelial senescence is also present in PAH PAECs and contributes to endothelial dysfunction and the progressive nature of pulmonary artery remodeling (5, 6). Our findings further show that RAB7 knockdown affected transcription of gene clusters affecting organ and tissue survival, cell death, cell growth and inflammatory, development and immune response. The angiogenic deficit and senescence also shows in the transcriptomic profile of RAB7 silenced PAECs. Because of its fundamental function in endosome and lysosome function and autophagy/mitophagy, we show in our work that endothelial RAB7 indeed has an important protective function in the pulmonary vasculature.

To determine if enhancing the activity of remaining RAB7 provides a therapeutic avenue, we used the GTPase activator ML-098 which has a low ED_50_ for RAB7, and much smaller affinity to other small GTPases (43). ML-098 treatment not only prevented, but also reduced established PH in rats exposed to Hx/Su. ML-098 has only been used in cell culture experiments, where authors have shown to protect from age-related deterioration of oocytes (19), which is consistent with our observation that loss of RAB7 promotes cell cycle arrest and senescence.

Our study has limitations: 1. Our work focused on the role of RAB7 in EC function and lung vascular remodeling but other aspects of RAB7 deficiency such as an exaggerated inflammatory response may also be mechanistically relevant. 2. We have tested ML-098 only in the Hx/Su model of PH, although this is one of the most relevant models mimicking many features of PAH, including the progressive nature of the condition and the occlusive pulmonary arteriopathy (22, 44).

Taken together, we demonstrate here in PAECs a fundamental role for the endolysosomal GTPase RAB7 in physiologic cell function, transcriptomic profile, endolysosome function, mitochondrial health, and pulmonary vascular integrity. Pharmacologic modulation of RAB7 function could complement existing therapeutic strategies and offer a new therapeutic avenue for patients with PAH.

## METHODS

### Reagents and constructs

#### Antibodies

α-SMA (M0851, DAKO), β-actin (127M4866V, Millipore Sigma), RAB7 (ab137029, Abcam), PCNA (25865, Cell Signaling Technologies), RAB7 (957465, Cell Signaling Technologies), vWF (A0082, Millipore Sigma).

#### Recombinant DNA and RNA

Ad-sh-RAB7 (2001, Vector Biolabs), Ad-sh-scrm (1122-HT, Vector Biolabs), RAB7 siRNA (NM_004637 13.2, IDT), control siRNA (51-01-14-04, IDT).

#### Other Reagents

7-AAD (51-68981E, BD Biosciences), Annexin V (APC) (550475), Annexin V binding buffer (51-66121E, BD Biosciences), BD Bioscience), Cathepsin B assay kit (PK-CA577-K140,), CellLight^™^ Early Endosomes-GFP BacMam 2.0 (C10586, Thermo Fisher), CellLight^™^ Mitochondria GFP BacMam 2.0 (C10600, Thermo Fisher), BrdU APC flow kit (51-9000019AK, BD Biosciences), DAPI (D1306, Invitrogen), Hoechst (H21486, Thermo Fisher), Lysotracker green (L7526, Thermo Fisher), ML-098 (T4619, TargetMol), MitoSOXTM (M36008, Thermo Fisher), Mito TrackerTM green (M7514, Thermo Fisher), pHrodoTM Red Dextran (P10361, Invitrogen), SU5416 (S8442, Sigma), Matrigel (356230, Corning), TMRE mitochondrial membrane staining kit with FCCP (ab113852, Abcam), TMRM Assay Kit (ab228569, Abcam).

### Tissue samples

De-identified Formalin-fixed, paraffin-embedded 5 μm tissue sections were obtained from the Pulmonary Hypertension Breakthrough Initiative (PHBI). De-identified human tissue samples were deemed as “non-human subjects research” from the Office of Research Subjects Protection at OSU.

### Animal experiments

RAB7^fl/wt^Cdh5-Cre mice were generated by crossbreeding RAB7^fl/fl^ mice (strain *B6.129(Cg)-Rab7^tm1.1Ale^*/J, #021589, Jackson Labs) with vascular endothelial cadherin (VE-cadherin, Cdh5) Cre mice (strain B6.Cg-Tg(Cdh5-cre)1Spe/J, #033055, Jackson Labs). The mice were crossbred for multiple generations before the experiments. Male and female mice aged 8-16 weeks were used for the experiments. Genotyping was performed by Transnetyx. Littermate RAB7^fl/wt^Cdh5-Cre^-^ mice were used as controls in the experiments. For rat experiments, Sprague Dawley rats (age 6 weeks) were obtained from Envigo. For the chronic hypoxia + SU5416 (Hx/Su) rat model, male rats were treated with SU5416 s.c. (20 mg/kg bodyweight) at the beginning of 3 weeks of chronic hypoxia exposure (inspiratory oxygen fraction 10%) in a normobaric nitrogen dilution chamber as published previously (45). For the Hx/Su mouse model, mice were given 20 mg/kg SU5416 s.c. once a week during a 3-week exposure to chronic hypoxia (45). ML-098 was diluted in dimethyl sulfoxide (DMSO) and given to male rats after dilution in 0.9% saline (final 1% DMSO) at 0.1, 1.0 and 10 mg/kg bodyweight five times a week from day 1-21 (preventive) during the chronic hypoxia phase of the cHx/Su model or day 22-35 after return to normoxia. For all treatments, animals were randomly assigned to the treatment groups and treatments were given in a blinded manner. At the indicated time points, echocardiography was performed with using a Vevo 2100 system (Visual Sonics, Toronto, ON, Canada) under isoflurane anesthesia (Small Animal Imaging Facility at OSU) or a GE Vivid IQ Premium system (GE Healthcare, Chicago, IL) under ketamine/xylazine anesthesia. Echocardiographic estimation of the cardiac output was calculated according to (46). Terminal right heart catheterization was done with a 1.4F Millar catheter and Powerlab acquisition system (AD Instruments, Colorado Springs, CO). For this procedure, the animals were anesthesized with ketamine/xylazine and ventilated after tracheostomy with a small animal ventilator (RoVent, Kent Scientific Corporation, Torrington, CT). A median sternotomy was used to open the chest cavity and a 1.4F Millar catheter was inserted following a small puncture of the right ventricle. Right ventricular hemodynamics were recorded over at least 5 min for steady state measurements. Acquisition and analysis of echocardiographic and hemodynamic data was blinded by numerical coding.

### Cell culture experiments

Human PAECs were cultured in complete EGM-2MV media (Lonza) in a cell culture incubator at 37°C with 5% CO2 and 100% humidity. For siRNA-mediated knockdown of RAB7, PAECs were transfected with 50 nM RAB7 or control siRNA using GenMute Reagent (SL100568 SignaGen). After 24h, siRNA was removed and experiments were performed at 48h or 72h. To test 2D angiogenic network formation, cells were seeded on Matrigel in μ-slides Angiogenesis (Ibidi) and images of network formation were acquired at 4× magnification using a EVOS M7000 automatic microscope. The images were analyzed using AngioTool software (NCI). To test gap closure, ECs were seeded in 3 well migration assay culture inserts. Removal of the insert left a well-defined uniform gap and gap closure was observed after 24h as published previously (45). For DNA synthesis and cell cycle analysis, cells were pulsed for 4h with 5-Bromodeoxyuridine (10 μM), then cells were removed using Accutase for flow cytometry staining using the APC BrdU flow kit (BD Biosciences). Analysis was performed using a BD FACSSymphony A1 flow cytometer and FlowJo 10 software (FlowJo). Data were prepared as cell cycle analysis and % BrdU^+^ cells. To detect cell proliferation, we seeded 50,000 cells in 6 well plates, followed by transfection with siRNA. After 72h, cells were removed, and cell count was measured using a TC20 Automated Cell Counter (BioRad). To identify apoptosis, cells were staining using the APC Annexin V staining kit (BD Bioscience) and the fraction of Annexin V^+^ 7-AAD^-^ cells was evaluated using FlowJo 10 software. To detect cellular senescence, cells were stained with senescence-associated (SA)-β-galactosidase (gal) according to the manufacturer’s instructions for Senescence β-Galactosidase Staining Kit (Cell signaling, 9860). Cells were counterstained with DAPI. SA-β-gal activity was determined by the detection of blue-green precipitate over the cells. Cells were viewed using EVOS M7000 automated fluorescence microscope (Invitrogen) under bright field. To evaluate vesicular trafficking and degradation, cells were incubated with pHrodo dextran (concentration) for 20 min, followed by fixation with 10% formalin. To specifically determine endosomal localization of dextran, GFP-conjugated RAB5A (early endosome marker) was expressed using a BacMam 2.0 baculovirus construct using the CellLight™ Early Endosomes-GFP kit. To obtain the lysosomal localization of dextran, lysosomes were stained with lysotracker green. To measure lysosome activity, a cathepsin B activity assay was used according to the manufacturer’s recommendations. To test mitochondrial membrane potential, we used the Tetramethylrhodamine ethyl ester (TMRE) staining kit (200 nM) combined with MitoTracker™ Green (mitogreen, 100 nM) staining. Staining was performed in fetal calf serum free media for 30 min. FCCP was added 10 min before staining at 20 μM. Analysis was performed using a BD FACSSymphony A1 flow cytometer and FlowJo 10 software (FlowJo). Data were presented as fraction of TMRE^high^ mitogreen^high^ (physiologic ΔΨ_m_) and TMRE^low^ mitogreen^high^ cells (decreased ΔΨ_m_). ΔΨ_m_ distribution was also determined using TMRM staining on adherent PAECs. To determine mitochondrial ROS production, cells were stained with 2 μM mitoSOX mitochondrial ROS staining for 20 min. combined with LIVE/DEAD™ Fixable Near-IR Dead Cell Stain Kit, followed by analysis on BD FACSSymphony A1 flow cytometer and FlowJo 10 software. Lactate production was measured with the Lactate-Glo™ Assay (Promega) according to the manufacturer’s recommendations.

### Determination of Mitochondrial Dynamics

Mitochondrial motility (distribution of net mitochondrial movement) was determined using wide-field fluorescence microscopy and image analysis as described in significant detail in our previous work (47). Briefly, the day prior to experiments, cells were transfected with CellLight Mitochondria-GFP, BacMam 2.0 (Life Technologies, Grand Island, NY, USA) at 40 particles/cell and kept under dark conditions at 37 °C. Cells were also stained with DAPI and were placed in Recording HBSS (HBSS pH 7.4 with 1.3 mM CaCl2, 0.9 mM MgCl2, 2 mM glutamine, 0.1 g/L heparin, 5.6 mM glucose, and 1% FBS) for microscopy imaging. For image acquisition, cells were imaged using an Olympus IX-51 inverted epifluorescence microscope with an Olympus 40x oil immersion objective lens, a Hamamatsu ORCA camera (2048 × 2048 pixels) with an LED light source from Lumencor. Images were collected using Metamorph 7.10.5 software. Images of cells were captured every 3 seconds over a total duration of 5 minutes, providing 101 sequential image frames which were referred to as “whole cell” frames. Image pre-processing followed by analysis of intracellular mitochondrial motility in the cell perinuclear and cell peripheral regions was performed using ImageJ and MATLAB with established methodology (48). This degree of analysis was performed to determine if intracellular inhomogeneities of mitochondrial motility were present and to quantify measureable effects of specific cellular conditions.

### Seahorse OCR/ECAR measurement

Cellular mitochondrial oxygen consumption rates (OCR) in pmol/min and extracellular acidification rates (ECAR) in mpH/min were evaluated using the Seahorse XFe24 Analyzer (Agilent, Santa Clara, CA). Experiments were performed with unbuffered DMEM XF assay media supplemented with 2 mM GlutaMAX, 1 mM sodium pyruvate, and 5 mM glucose (OCR experiments only) at pH 7.4 and 37 °C. Following instrument calibration, select compounds were injected to obtain basal respiration, LEAK and maximum respiration (Max) in OCR experiments. After initial measurement of basal respiration, oligomycin (2 μM) was injected to inhibit ATP synthase (Complex V or CV) to obtain LEAK. FCCP (2.5 μM) was subsequently injected to induce uncoupling that yields Max. We also obtained spare respiratory capacity (SRC), which assesses mitochondrial reserve for ATP production in response to increased metabolic demand. SRC is equal to the difference between maximal and basal respiration. In the ECAR experiments, sequential administration of glucose (10 mM) and oligomycin (1 μM) enables measurement of glycolysis and glycolytic capacity, respectively. Subsequent injection of 2-deoxy glucose (50 mM) enables calculation of glycolytic reserve and non-glycolytic acidification.

These respiratory states and parameters of glycolytic function were measured at consecutive time points, which were averaged to give a single value for each state for each cell culture well. Data were normalized using metabolic activity; background readings for each plate were calculated by averaging the OCRs/ECARs from the background wells for those plates and were subtracted from all subsequent readings. Erratic or extreme background well readings were excluded. All experiments were repeated in duplicate in runs performed on separate days. Each well was analyzed individually, and the resultant data are presented as the mean ± standard deviation (SD) for all experiments.

### Protein isolation and immunoblotting

Protein isolation and Western blotting were performed as published previously (4, 45, 49). Following lysis with Radio immunoprecipitation buffer (RIPA) with proteinase and phosphatase inhibitors, 10-20 μg protein (cell lysate) and 40 μg protein (whole lung lysate) was resolved by SDSPAGE at 90 V for 60-80 min. Then, protein was blotted at 270 mA for 2 hours onto a nitrocellulose membrane for antibody staining and chemiluminiscence detection (ECL). Membranes were blocked with 5% bovine serum albumin/0.5% Tween20/TBS for 1 hour. Primary antibody incubation was performed in 5% bovine serum albumin/0.5% Tween20/TBS overnight at 4 °C, followed by secondary antibody incubation in 5% bovine serum albumin/0.5% Tween20/TBS for 1h (horse radish peroxidase conjugated secondary antibodies). Images were obtained by developing with ECL solution and image acquisition with a Biorad ChemiDoc gel imager with ImageLab software (Biorad, Hercules, CA). Densitometry with automated background substraction was performed with ImageLab (Biorad, Hercules, CA).

### RNA isolation and real-time quantitative PCR

RNA isolation, cDNA preparation and quantitative real-time PCR were performed as published previously (4, 45, 49). RNA was isolated using the miRNAeasy kit (Qiagen) according to the manufacturer’s instructions. For reverse transcriptase (RT) reaction, DNAase I treatment was performed at 0.1 U/μl final activity for 15 min at room temperature, followed by addition of 25 mM EDTA to inactive DNase and incubation at 65 °C for 10 min. For the reverse transcriptase (RT) reaction, a mastermix was prepared of 10× RT buffer, dNTP mix, RT random primers and RNAse free water and added to the RNA samples. RT reaction was run with the following steps: 10 min at 25 °C, 120 min at 37 °C and 5 min at 85 °C in a Thermocycler (T100, BioRad). For real-time amplification, cDNA samples were diluted 1:10 in nuclear free water. A mastermix was prepared for each well (384 well plate) consisting of 1 μl of forward and reverse primer mix, 5 μl SYBR Green mastermix (iTaq, Biorad) and 4 μl cDNA. QRT-PCR was run in a CFX384 touch qRT-PCR system (Biorad) with the following program: initial step at 95.0°C for 30 s. Then, 45 cycles of 95.0°C for 5 s and 60.0°C for 30 s. After a final step of 95°C for 10s, a melting curve was obtained by raising the temperature from 65.0 to 95.0°C in 0.5°C increments/5 s. The analysis was performed by using the method published in (50)

### RNA sequencing and analysis

RNA was isolated using the Qiagen miRNAeasy mini kit according to the manufacturer instructions followed by DNAse digestion to remove potential genomic DNA contamination. Total mRNA concentrations were measured using Nanodrop. RNA sequencing was performed in the Genomic Shared Resource at The Ohio State University using NEBNext^®^ Ultra^™^ II Directional RNA Library Prep Kit for Illumina, NEBNext rRNA Depletion Kit, NEBNext Multiplex Oligos for Illumina Unique Dual Index Primer Pairs. 100 ng total RNA (quantified using Qubit Fluorometer) was used as library input. Fragmentation time was 10 min and PCR was done 11x. Libraries were sequenced with Novaseq SP Pair_end 150 bp format.

Raw Data was analyzed by ROSALIND^®^ (https://rosalind.bio/), with a HyperScale architecture developed by ROSALIND, Inc. (San Diego, CA). Reads were trimmed using cutadapt (51). Quality scores were assessed using FastQC (52). Reads were aligned to the Homo sapiens genome build GRCh38 using STAR (53). Individual sample reads were quantified using HTseq (54) and normalized via Relative Log Expression (RLE) using DESeq2 R library (55). Read Distribution percentages, violin plots, identity heatmaps, and sample MDS plots were generated as part of the QC step using RSeQC (56). DEseq2 was also used to calculate fold changes and p-values and perform optional covariate correction. Clustering of genes for the final heatmap of differentially expressed genes was done using the PAM (Partitioning Around Medoids) method using the fpc R library (57). Hypergeometric distribution was used to analyze the enrichment of pathways, gene ontology, domain structure, and other ontologies. The topGO R library (58) was used to determine local similarities and dependencies between GO terms in order to perform Elim pruning correction. Several database sources were referenced for enrichment analysis, including Interpro (59), NCBI (60), MSigDB (61, 62), REACTOME (63), WikiPathways (64). Enrichment was calculated relative to a set of background genes relevant for the experiment. Following identification of differentially expressed genes with fold changes of >1.25 and an adjusted P value of <0.05, downstream analysis of functions and pathway enrichment was performed using Ingenuity Pathway analysis (Qiagen).

### Histology and immunohistochemistry

Immunohistochemistry and double immunofluorescence stainings were performed according to previous publications (4, 45). In brief, for immunohistochemistry, rehydration of slides was followed by antigen retrieval (20 min. heating in boiling citrate buffer at pH 6.0 in most protocols, except for vWF, which used proteinase K 1:50 for 5 min as antigen retrieval). Blocking of endogenous peroxidase (H2O2) and of unspecific binding with 1% normal swine serum (NSS)/PBS for 15 min was followed by incubation with primary antibodies (diluted in 1% NSS/PBS) overnight at 4°C. Further steps were incubation with secondary biotin-labeled antibody (1 hour, diluted in 1% NSS/PBS) and incubation with HRP conjugated Streptavidin for 1 hour. Slides were stained with diaminobenzidine solution and counterstained with Mayer’s hematoxylin. After dehydration, slides were mounted with coverslip and permanent mounting medium.

Immunofluorescence required rehydration followed by antigen retrieval (20 min. heating in boiling citrate buffer pH 6.0). After blocking unspecific binding with 1% normal swine serum (NSS)/PBS for 15 min, primary antibodies were added 1% NSS/PBS for overnight incubation at 4°C. Incubation with secondary fluorescence-labeled antibody (1 hour, diluted in 1% NSS/PBS) was followed by nuclear staining with DAPI for 5 min. Then, slides were mounted with coverslip and SlowFade Gold.

Media wall thickness (MWT) was analyzed as described previously (4, 49). In brief, images of pulmonary arteries were acquired with an EVOS M7000 automated microscope. Treatment groups were masked by numerical coding. Media thickness (MT) and external diameter (ED) were measured and MWT was calculated as MWT=[(2×MT)/ED]×100%. Pulmonary arteries were categorized as small-sized 25 μm < ED < 50 m and medium-sized 50 μm ≤ ED < 100 μm. For each animal, 30-40 pulmonary arteries were measured in two orthogonal directions using Fiji image analysis software. Pulmonary artery occlusion in rats was analyzed as published previously by us using slides stained for vWF immunohistochemistry and identifying the fraction of small-sized pulmonary arteries that were classified as ‘patent’, ‘partially occluded’ and ‘completely obstructed’ (4, 45).

For analysis of PCNA^+^ cells per pulmonary artery, images of pulmonary arteries were randomly acquired from transversal sections stained for PCNA immunohistochemistry at 40× magnification. The number of PCNA^+^ cells was divided by the number of nuclei/pulmonary artery wall crossection to calculate the fraction of PCNA^+^ cells.

For all histomorphometric analyses, objectivity was ensured by masking treatment groups through numerical coding and an investigator unaware of the treatment groups.

### Microscopy

Light microscopy images were acquired using an EVOS M7000 automated microscope (Thermo Fisher Scientific, Waltham, MA). Immunofluorescence images were acquired using an inverted Olympus FV3000 confocal microscope system located at the OSU Campus Microscopy and Imaging Facility. The images were assembled with Fiji software.

### Statistical analysis

Data were compared using Student’s t-test (2 groups), or one or two-way ANOVA (more than 2 groups), following by multiple comparison correction using the Holm-Sidak or Sidak tests. Normal distribution was tested with the Shapiro-Wilk test. The calculations were performed using Prism 9.0 (GraphPad Software Inc., San Diego, CA). A *P* value of <0.05 was considered significant.

### Study approval

De-identified human tissue samples and cell lines were deemed as “non-human subjects research” from the Office of Research Subjects Protection at OSU. Animal experiments were approved by the IACUC at OSU under protocol number 2019A00000092.

## ACKNOWLEDGEMENTS

The authors acknowledge the expert technical help of Dr. Mehboob Ali, Kyle Shin, Alexander Pan, Jaylen Hudson, and Pranav Gunturu. The study was supported by grants from the NIH/NHLBI (HL139881) to L.F., (HL130261, HL113178, HL159638, and HL103455) to E.A.G. The study was further supported by a grant from the Office of Naval Research Grant (ONR) N000142212170 to D.M.E. Data/Tissue samples provided by PHBI under the Pulmonary Hypertension Breakthrough Initiative (PHBI). Funding for the PHBI is provided under an NHLBI R24 grant, R24HL123767, and by the Cardiovascular Medical Research and Education Fund (CMREF). The authors acknowledge resources for confocal microscopy from the Campus Microscopy and Imaging Facility (CMIF), and the OSU Comprehensive Cancer Center (OSUCCC) Microscopy Shared Resource (MSR) at the Ohio State University with NIH S10 OD025008 and NIH NIC P30CA016058. Echocardiography was, in part, performed in the Small Animal Imaging Core at OSU. These facilities are supported in part by grant P30 CA016058, National Cancer Institute, Bethesda, MD. RNA sequencing was performed in the Genomic Shared Resource of the OSU Comprehensive Cancer Center, partially supported by grant P30-CA016058 from the National Cancer Institute. The content is solely the responsibility of the authors and does not necessarily represent the official views of the National Institutes of Health.

## AUTHOR CONTRIBUTIONS

B.P., S.B., N.S., A.R.B., H.Y., R.K.M., E.A.G., D.M.E. and L.F. conceived and designed the study; B.P., S.B., T.H., D.F., L.R., A.G., G.N., A.R.B., T.K., P.Y., D.E.M. and L.F. performed experiments, obtained and analyzed data; B.P., S.B., D.M.E. and L.F. wrote the initial draft of the manuscript; all authors contributed to manuscript revision; all authors approved the final version; the order of co-first authors was based on the overall contribution to data acquisition and manuscript preparation.

